# ASAREE: An Analytical Sandbox for Agentic AI Research, Engineering, and Experimentation

**DOI:** 10.64898/2026.08.20.746074

**Authors:** Jay Moran, Philip J. Freda, Attri Ghosh, Corey T. Walker, Miguel E. Hernandez, Jason H. Moore

## Abstract

**Summary:** Agentic AI platforms enable the engineering of autonomous workflows but are not designed for experimentation and hypothesis testing. **ASAREE** (Analytical Sandbox for Agentic AI Research, Engineering, and Experimentation), is an open-source platform to address this gap. ASAREE creates agents, connects to MCP servers and tools, and designs factorial experiments through a visual interface or Python SDK. It records a full provenance trace for every run and routes all model calls through a provider-agnostic bridge that supports local deployments, ensuring data privacy. As a use-case, we use ASAREE to evaluate key design choices in a mutli-agent machine learning pipeline. Across a 2 *×* 2 *×* 2 factorial design, more advanced models, greater reasoning effort, and critic agent use significantly increased compute time, token use, cost, and feature count without improving predictive performance. The lowest-cost baseline, Claude Sonnet 5 with medium effort and no critic, achieved the highest mean PR AUC while Claude Opus 5 with extra high effort and a critic agent cost 15.5*×* more (USD) and ran 13.1*×* longer while performing worse on average. These findings highlight ASAREE as a robust framework for evaluating agentic system performance and resource efficiency.

**Availability and implementation:** ASAREE is available on GitHub at: https://github.com/EpistasisLab/ASAREE.

**Supplementary Information:** Supplementary information is available at https://github.com/EpistasisLab/ASAREE/tree/main/publications/bioinformatics

## 1 Introduction

Agentic AI systems that reason goals, call external tools, use specialized skills, access curated knowledge, and act over multiple steps are increasingly being used across science and industry. In biomedicine, agents are being applied to hypothesis generation, literature synthesis, knowledge reasoning, and automation of data-analysis pipelines [Chen et al., 2026, Zack et al., 2026]. The appeal is that agents can orchestrate the same heterogeneous tools scientists already use, such as databases, statistical packages, and predictive modeling approaches, all via natural-language.

This promise rests on a methodological gap. The performance of agentic systems depend on many design choices, including the underlying model, reasoning effort, whether decisions are critiqued, and how work is divided (the agent topology). Currently, these choices are rarely tested under controlled conditions or reported in enough detail to reproduce. General-purpose frameworks such as LangGraph and CrewAI are engineering toolkits for building agents rather than platforms for evaluating them. In other words, they are not designed to treat architectural decisions as experimental variables, to record step-level provenance in a form comparable across runs, or to enforce data governance constraints of sensitive biomedical data [LangChain, Inc., 2024, crewAI, Inc., 2024]. Domain-specific biomedical agents sit at the other extreme, typically single-purpose and not built for systematic comparison [Wang et al., 2025, Bran et al., 2024]. What is missing is a platform where researchers can both construct agentic workflows and rigorously evaluate their performance and reproducibility across model and architecture choices.

We present **ASAREE** (Analytical Sandbox for Agentic AI Research, Engineering, and Experimentation), a transparent, open-source platform addressing this gap. ASAREE executes agents through an explicit Sense-Reason-Plan-Act (SRPA) loop, routes tool calls through the Model Context Protocol (MCP), and records results for auditing and reproducibility. Specifying agentic architectures as configurations rather than custom code lets competing designs be compared under identical conditions. Additionally, ASAREE is not tied to any specific model provider and workflows can run entirely on local/secure models so data privacy can be maintained.

## 2 Implementation

ASAREE is an open-source application built on top of Motoro, an open-source, pinned library dependency which provides agent execution primitives. ASAREE adds an HTTP API, auth, persistence, and a node-based UI using React Flow.

### 2.1 ASAREE Architecture

1. **The SRPA loop**. Every agent turn is an iteration of four explicit phases. Sense gathers inputs (the task, prior memory, and the available tools), Reason interprets inputs against the agent’s goal to produce a strategy, Plan decomposes that strategy into ordered steps, and Act executes the steps. Reason and Plan are deliberately separated into distinct model calls with distinct outputs, so that each can be inspected and evaluated independently. Each phase is recorded as a step, including its inputs, outputs, model call details (tokens, latency, and cost), and any tool calls. Every completed run also includes an structured output envelope, which is a uniform contract (status, result, summary, artifacts, confidence, caveats) with an optional typed payload. Thus, a run’s result is read the same way regardless of the agent that produced it. Together, these yield a complete, auditable provenance trace for every run, which serve as prerequisites for reproducible scientific use.
2. **MCP as the tool boundary**. ASAREE does not hard-code data sources or analysis tools. It allows connection of MCP servers: the available tools are surfaced during Sense and invoked during Act, over either stdio (local servers) or streamable HTTP (remote servers). Any dataset, database, knowledge graph, or analysis routine is wrapped once as an MCP server that is then reused without modification across agents and experiments. This boundary is what makes an analytical workflows portable across problems: the same agent logic can be retargeted to a new dataset by pointing to different MCP server, with no change to the agent itself.
3. **Experimental design** Researchers can build an experiment by creating AI Agents and connecting components such as MCP Servers, Tools, and Datasets. This can be done either visually on the Frontend or via a Software Development Kit (SDK). Experimental design factors can be selected (*e*.*g*., factorial designs) and multiple levels can be configured. ASAREE generates a design and the cells for the experiment. Performance metrics can be added for downstream reporting and analysis.
4. **Graphical User Interface (GUI)** ASAREE ships with a Graphical User Interface (Figure 1A) based on React Flow where experiments can be fully designed. Components (Agents, LLM provider, MCP Tools, Architectural Patterns) are represented as nodes in a graph configured through an interactive canvas. Researchers have the option to download their experiment in JSON format, which can be shared and imported into the same or other instances of ASAREE for reproducibility.
5. **Software Development Kit (SDK)** Advanced users can use the SDK. ASAREE ships with a Python client named ‘asaree-client’. The asaree-client provides all the capabilities needed to fully create and run experiments programmatically. This makes ASAREE extensible. Although only a Python SDK client exists today, SDK clients for other programming languages can be easily built by the open-source community.
6. **Provider independence**. All model calls go through a single bridge built on litellm and Instructor, so agents are not tied to any one vendor. Cloud providers (*e*.*g*., Anthropic, OpenAI) and locally hosted models served behind an OpenAI-compatible endpoint are addressed through the same interface and structured-output contract. This allows analyses to run entirely on local models - important when data governance precludes sending sensitive biomedical data to a third-party API. Also, it allows models to be compared head-to-head under identical agent configurations.
7. **Deployment** Instructions to run ASAREE are available on the GitHub repository. It is deployed through docker containers for portability and once the application is deployed and running, users can access the Frontend on their web browser. Registered users can then generate their own API key for asaree-client to use the SDK.

**Fig. 1.**
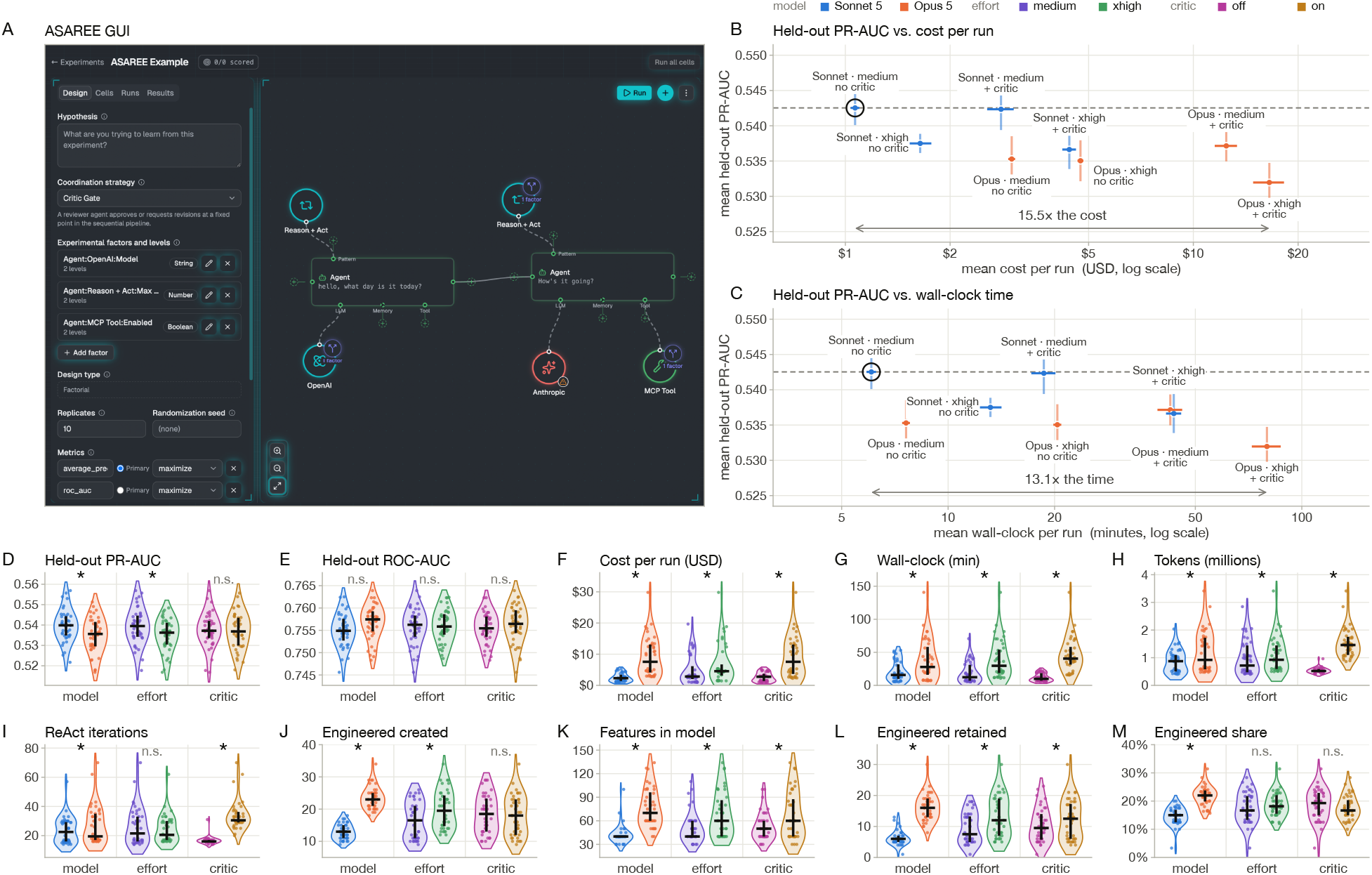
ASAREE GUI and Use Case Results. (A) Screenshot of an example ASAREE experiment. (B and C) Pareto fronts of mean held-out PR AUC across configurations v. mean cost and mean wall-clock time (m) in log scales. Bars are *±*1 SE. Circled points denote Pareto-optimality. (D-M) Violin plots of each endpoint at low and high levels of each factor. Left violins represent baseline levels (Sonnet 5, medium effort, critic off). Endpoints are accuracy (D, E), resource use (F-I), and feature-set compositions (J-M). Asterisks denote significant main effects after Holm adjustment, *n.s*. otherwise.

## 3 Case Study

### 3.1 Data Description

Our data consists of an IRB-approved retrospective cohort of 7,108 elective spinal fusion surgeries performed at Cedars-Sinai Medical Center in Los Angeles from 2013 to 2023. Patients aged 18-85 and with *≥* 2 spinal fusion surgeries. The dataset contains 126 predictors and a binary outcome, non-home discharge (27.3% rate), defined as post-surgical discharge to any destination other than the patient’s home.

Predictors include demographics, social history, pain scores, past medical history, medications, and laboratory values. Most features are derived from our prior work on this cohort [Ghosh et al., 2025, 2026]. Additional features are intentionally added to evaluate agent performance in realistic clinical data-integrity challenges, including correlated variables, high missingness, and clinically implausible values.

### 3.2 Multi-Agent System Architecture and Evaluation

We develop four agents with role-associated prompts, tools, and goals: data cleaning (DC); feature transformation/engineering (FTE); feature selection (FS); and machine learning/modeling (MLM). The agents run as a linear pipeline in that order (1 turn per agent), each passing a modified training dataset and modification recipe to the next agent.

The DC agent identifies and removes outliers, imputes missing values, and deletes low variance columns before passing the data to the FTE agent, which makes feature engineering decisions on the cleaned dataset. The FS agent removes features with low association with the training outcome. Finally, the MLM agent specifies nine XGBoost [Chen and Guestrin, 2016] hyperparameters controlling capacity, learning rate, complexity, subsampling, and regularization. These are passed and executed by a fixed script rather than by an agent to ensure run completion. No agent ever observes the test set.

Every decision an agent makes is recorded in the modification recipe, which is replayed on the test set at evaluation time to mirror training modifications without refitting.

We evaluate fitted models on the transformed test set using ROC AUC and PR AUC. Additionally, we record total tokens used, US dollars spent (USD), wall-clock time (minutes), and agent ReAct iterations. ReAct, or reason-act-observe cycles, are SRPA instances where the agent reasons about the current state, calls a tool, and observes the result before deciding whether to iterate again or finish its turn [Yao et al., 2022]. In other words, one or more SPRA iterations occur in a ReAct cycle. We also record the number of features constructed by the FTE agent, the size of the final feature set, and the number and proportion of engineered features in the final set.

### 3.3 Experimental Design and Statistical Analysis

We construct a 2 *×* 2 *×* 2 factorial design to evaluate the effects of model, reasoning effort, and a critic agent. The model factor compares Claude Sonnet 5 and Claude Opus 5. For effort, we compare medium and extra high (xhigh) effort. Anthropic’s effort is an API-level control over adaptive thinking that governs how readily the model spends tokens on reasoning, tool usage, and response [Anthropic, 2026]. The critic factor toggles a fifth agent that reviews the modification recipe of each agent at the end of its turn. The critic either approves the recipe, advancing the pipeline, or requires the agent to run one additional turn to add, remove, or revise recipe decisions. The critic does not re-evaluate the second turn. The critic utilizes the model and effort setting of the current factorial cell. Ten replicates of the eight cells are run on a fixed 70/30 train/test split (Seed 42), so that run-to-run variability can be separated from between-cell differences.

Models are prespecified according to the distribution of the outcome. Beta regressions with logit links [Ferrari and Cribari-Neto, 2004] are used for ROC AUC and PR AUC. USD cost, wall-clock time, and count outcomes (tokens, ReAct iterations, and feature counts) use log links, with gamma regression for continuous outcomes and count models for counts. Negative binomial regression is used for overdispersed counts, and Poisson regression with Heteroskedasticity-Consistent type 0 robust standard errors otherwise [White, 1980]. The proportion of engineered features in the final set is modeled with a binomial generalized linear model, using final set size as the denominator. Main effects are estimated from the eight predicted cell means and reported on the response scale with delta method 95% Wald intervals, with effects expressed as differences for logit-link models and ratios for log-link models. The model effect on PR AUC is the primary test at (*α* = 0.05). Effort and critic effects on PR AUC are Holm-adjusted as a family of two. For each remaining endpoint, the three main effects are Holm-adjusted within that endpoint. Interactions are tested using the corresponding link-scale coefficient, rather than an eight-cell response-scale contrast, which is not cleanly separable from main effects under a log link. Therefore, we report interaction confidence intervals and unadjusted *p*-values without error-rate control. Finally, the eight configurations are compared for each endpoint using Kruskal-Wallis tests followed by Holm-adjusted pairwise Mann-Whitney tests. Full specifications, analyses, and code are available in GitHub.

### 3.4 Results and Discussion

The full experimental pipeline used $466.23 and 38.6 hours of compute. Four fifths of this cost was attributable to Opus 5, which predicted no better than Sonnet 5. Overall, more resource use, including higher effort and the critic agent, led to no gain in predictive performance (Figure 1B-M).

#### 3.4.1 Accuracy

Sonnet 5 performs slightly better than Opus 5 on held-out PR AUC (0.5397 v. 0.5349, *p* = 0.0056), while medium effort performs slightly better than xhigh effort (0.5393 v. 0.5353, Holm-adjusted *p* = 0.044). The critic has no measurable effect (*p* = 0.75). Although Sonnet 5 and medium-effort improvements are consistent, they are small relative to the variation across runs. None of the individual configurations are different from each other after adjustment. There are no significant ROC AUC effects after adjustment.

#### 3.4.2 Cost and Feature Selection/Engineering

The critic is the most expensive factor. It increases wall-clock time by 3.90*×*, USD cost by 3.46*×*, token use by 2.86*×*, and ReAct iterations by 2.03*×*. Increasing effort doubles wall-clock time and raises cost by 41%. However, xhigh effort does not significantly change the number of ReAct iterations (*p* = 0.22), suggesting that extra computation happens within iterations. Opus 5 costs 3.71*×* more USD than Sonnet 5 while also producing substantially more engineered features: it creates 1.82*×* as many, retains 2.59*×* as many after selection, and increases their share of the final set from 14.6% to 21.8% (*p* = 2.3 *×* 10^*−*25^), a 1.50*×* relative increase. Opus also produced larger feature sets. Yet, none of this, nor the critic agent translated into better held-out performance (Figure 1B-E).

#### 3.4.3 Interactions

Opus 5 with critic increases resource use more than either factor alone and is observed for wall-clock time (1.47*×*), token use (1.39*×*), and USD cost (1.43*×*). These are the only three interactions (out of 44) that remain significant after correction.

#### 3.4.4 Cost–Accuracy Tradeoff

Sonnet 5 with medium effort and no critic agent also has the highest mean PR AUC, at $1.07 and 6.1 minutes per run on average. Opus 5 with xhigh effort and a critic has the lowest mean PR AUC, costing $16.53 and taking 79.7 minutes per run on average. It therefore costs 15.5*×* more and takes 13.1*×* longer to achieve worse performance.

## 4 Conclusions

ASAREE provides a transparent framework for systematically evaluating how design choices affect the behavior, resource use, and performance of agentic pipelines. Advanced models, higher effort, and a critic agent substantially increase costs without improving predictive performance. Although these results are specific to one dataset and prediction task, these results highlight the importance of evaluating agentic pipelines on both held-out performance and resource use rather than assuming that greater computational effort translates into better results.

## Acknowledgments

This work is supported in part by funds from the Center for AI Research and Education (CAIRE) at Cedars-Sinai Medical Center and grants from the National Institutes of Health USA (U01 AG066833, R01 LM014572, and P30 AG094848 awarded to JHM and K01 DA063751 awarded to PJF). The authors thank Mr. Nicholas Matsumoto for many fruitful discussions about agent engineering.

